# Complex Indel Detection: A Simulation-Based Framework and Parsing with FreeBayes

**DOI:** 10.64898/2026.05.26.727999

**Authors:** Yong Hwee Eddie Loh, Michael R. Lieber, Chih-Lin Hsieh, Zarko Manojlovic

## Abstract

In contrast to simple deletions and simple insertions, most complex indels involve both deletions and insertions, often with base changes within a few nucleotides of the indel’s left and right boundaries. These complex indels often arise from double-strand breaks (DSB), which in normal somatic cells are predominantly repaired by nonhomologous DNA end joining (NHEJ). Such complex indels pose a difficult analytical problem for existing indel callers because the observed VCF representation may be locally shifted, extended with matching flanking bases, or fragmented into several closely spaced calls. To evaluate complex indel representation, we tested six variant calling approaches: FreeBayes, HaplotypeCaller, Mutect2, Strelka2, DRAGEN Germline, and DRAGEN Somatic pipelines. Among the approaches evaluated, FreeBayes most consistently represented simulated complex indels as single nearby variant records. We then developed a parsing workflow that derives effective deleted and inserted sequences from FreeBayes VCF output and enriches for candidate complex indels. This approach supports analysis of naturally occurring DSB repair events in single human colon crypts.

## INTRODUCTION

Small sequence variants identified from short-read whole-genome sequencing are commonly summarized as single-nucleotide variants and simple insertions or deletions^1–3^. However, representation of variants involving both sequence deletion and sequence insertion at the same site is complicated by nearby mismatched bases or local sequence contexts that permit more than one valid representation of the same event^4^. As a result, a single biological alteration may be reported as one compound variant, split into several neighboring variants, or represented with additional matching flanking bases in the REF and ALT alleles. This ambiguity complicates detection, comparison across methods, and downstream biological interpretation.

This issue is especially relevant for DNA double-strand break (DSB) repair, predominantly repaired by nonhomologous end joining (NHEJ) and, less commonly, by alternative end joining (aEJ)^5^. NHEJ is the major form of repair at DSB sites in most vertebrate somatic cells, particularly in all phases of the cell cycle outside of S phase^5,6^. Based on biochemical and cellular studies, NHEJ repair products often include a short deletion together with nucleotide addition or a nearby base change at the repair junction^6^. In this manuscript, such repair products are referred to as complex indels because simple deletion, simple insertion, or isolated substitution does not fully describe these events. When deleted and inserted sequences share local similarity, the apparent junction can shift, the event can be left or right aligned in different ways, and the same repair product may appear either as a single compound event or as several smaller independent events^7^.

This problem became evident during our recent study of naturally occurring DNA damage and NHEJ repair in normal human colon crypts^5,8^. In that setting, complex repair associated variants were biologically expected, yet they were not readily apparent among routine small indel outputs from commonly used somatic callers such as Mutect2^9,10^ and Strelka2^11,12^. This raised the possibility that relevant repair events were present in the sequencing data but were not represented in a form that preserved their full, complex structure. These observations provided the practical motivation for developing a framework to evaluate complex indel representations and a downstream method suited to their interpretation.

The present study examined how commonly used variant callers handle complex indels and developed an approach to interpret FreeBayes output for these events. Complex indels were introduced *in silico* across a defined region of human chromosome 22, followed by simulated paired-end sequencing, alignment, and comparative variant calling with FreeBayes^13^, GATK HaplotypeCaller^14^, Mutect2^9,10^, Strelka2^11,12^, DRAGEN Germline (Illumina, Inc.)^15^, and DRAGEN Somatic (Illumina, Inc.)^16^. We determined that open-source FreeBayes was the most capable caller among the workflows tested for representing simulated complex indels as single nearby variant records. A parsing workflow was then developed to derive the effective deleted and inserted sequences from FreeBayes variant records for downstream complex indel analysis in the single colon crypt dataset.

## MATERIALS AND METHODS

### Code availability

Custom scripts used for complex indel simulation and FreeBayes parsing are available in the GitHub repository at https://github.com/eddieloh-usc/ComplexIndelSim.

### Software and custom scripts

The simulation and analysis workflow used the ART^17^ read simulator to generate synthetic Illumina paired-end reads with an available empirical Illumina error model, BWA MEM^18,19^ to align reads, SAMtools^20^ for BAM sorting and indexing, BEDtools^21^ for genomic interval operations, and six variant callers: FreeBayes, GATK HaplotypeCaller, GATK Mutect2, Strelka2, DRAGEN Germline (Illumina, Inc.), and DRAGEN Somatic (Illumina, Inc.). Two custom Perl scripts were used. The script, makeVariantFasta.pl generated simulated variant and reference FASTA files, along with a table of simulated events. The script, parseFreebayes.pl, processed FreeBayes VCF output and retained candidate complex indels after allele parsing and rule-based filtering.

### Generation of simulated complex indels

The Perl script, makeVariantFasta.pl was used to generate simulated complex indels within a selected interval of chromosome 22 from the GRCh38 reference genome. The extracted region spanned positions 30,000,001 through 31,000,000. Complex indels were introduced every 500 nucleotides. At each simulated site, 0 to 8 nucleotides from the reference sequence were deleted, and 0 to 8 random nucleotides were inserted. This design permitted a spectrum of observable events to be generated. These events range from pure insertions to pure deletions, as well as intermediate compound events reflecting the outcomes of sequence deletion followed by insertion. A fixed random seed of 123 was used for reproducibility. This procedure generated 2000 simulated variant sites. Three output files were produced, including a FASTA file containing the modified sequence, a FASTA file containing the unchanged reference interval, and a tab delimited table of simulated events.

### Simulation of sequencing reads

Synthetic Illumina sequencing reads were generated using art_illumina from both the altered sequence and the reference sequence. Reads were simulated as 150 bp paired-end fragments using the HS25 error model at 30-fold coverage, with a mean fragment size of 200 bp and a standard deviation of 10 bp. Separate FASTQ files were generated for the altered and reference sequences and were used as input for downstream alignment and variant calling.

### Alignment of simulated reads

Simulated reads from the altered and unchanged samples were aligned to the GRCh38 chromosome 22 reference sequence using BWA MEM. Read group information was added during alignment to ensure compatibility with downstream variant calling tools. Sorted BAM files were generated with SAMtools and indexed prior to variant calling. The altered sequence was treated as the variant sample, and the unchanged sequence was treated as the matched reference sample for callers requiring paired analysis.

### Variant calling

Six variant callers were evaluated. FreeBayes, HaplotypeCaller, and DRAGEN Germline pipeline were applied as germline callers. Mutect2, Strelka2, and DRAGEN Somatic pipeline were applied in paired tumor-normal mode, with the altered simulated sample designated as the tumor-equivalent sample and the unchanged simulated sample designated as the normal-equivalent control. FreeBayes was run jointly on both BAM files using default parameters, generating a single VCF for both samples. HaplotypeCaller was run separately on each sample in GVCF mode, followed by CombineGVCFs and GenotypeGVCFs to generate a final joint VCF. Mutect2 and Strelka2 were run using the altered sample BAM as the tumor and the unchanged sample BAM as the normal. DRAGEN Germline v4.5.4000 and DRAGEN Somatic v4.5.4000 pipelines were run using the Illumina DRAGEN standard pipeline. For DRAGEN Somatic, records tagged as mnv_component were excluded before counting nearby calls because these records represent component variants associated with combined multi-nucleotide variant records and would otherwise introduce redundant local calls. Strelka2 produces separate VCF files for SNVs and indels; these outputs were considered together when assessing nearby calls.

### Preliminary comparison of variant representation

Output from each variant caller was compared against the simulated event table by assessing whether a reported variant occurred within 5 bases of each simulated complex indel position. This window was used to allow for local representation differences arising from alignment and caller specific allele encoding. For the initial comparison, the number of reported variants within 5 bases of each simulated site was recorded in order to determine whether each simulated complex indel was represented as a single nearby event or as multiple nearby events. This comparison was intended to evaluate the representation of complex indels rather than merely positional recovery.

### Comparison of simulated and FreeBayes alleles

For simulated complex indels represented by a single nearby FreeBayes variant, the simulated deleted and inserted sequence was compared against the reported FreeBayes POS, REF, and ALT fields, where REF and ALT would reflect the predicted deletion and insertion sequence, respectively. A merged comparison file was generated to examine side by side correspondence between the simulated event and the called alleles. These comparisons showed that FreeBayes often reported additional matching flanking bases shared between REF and ALT. Accordingly, the interpretation focused on establishing a FreeBayes-derived prediction of effective deleted and inserted sequences, defined as the nonshared sequence remaining after removal of identical prefix and suffix bases from the reported REF and ALT alleles. In cases where the simulated deleted and inserted sequence themselves contained coincidental matching bases, the effective simulated complex indel rather than the exact original representation was also regarded as the relevant observable event. For sequence-level concordance analysis, one-to-one FreeBayes calls were compared with the simulated deletion and insertion before and after trimming the shared terminal sequence.

### Development of the FreeBayes parsing workflow

A custom Perl script, parseFreebayes.pl, was used to identify candidate complex indels from FreeBayes output. The complete rule-based workflow has been described previously by Loh YHE et al.^4^. In brief, the script excluded common and low complexity events, derived effective inserted and deleted sequences by trimming shared sequence from REF and ALT, and applied depth, allele frequency, repeat overlap, and sequence context filters to retain candidate complex indels.

### Parameters used in the parsing script

The parseFreebayes.pl script accepts the input VCF, output file, tumor sample name, control sample name, optional output file for discarded variants, minimum total depth, maximum control alternate allele frequency, minimum tumor alternate allele frequency, net insertion minus deletion threshold, repeat BED file, and reference FASTA file. In the simulation example, the script was applied to the FreeBayes VCF using the altered sequence as the tumor and the unchanged sequence as the control. The simulation-based application was used to test workflow behavior, while acknowledging that several filtering criteria were designed for experimental data rather than for the simplified simulation setting.

## RESULTS

### Standard somatic variant output underrepresented complex indels in colon crypt data

The motivation for the present framework arose from a prior analysis of naturally occurring somatic insertions in 106 single colon crypts with matched bulk controls, in which 16,483 autosomal insertion variants were identified from shared Mutect2 and Strelka2 output and then filtered to 77 high-confidence DNA breakage-associated insertion events^8^. Among those published 77 events, only seven were confirmed as complex indels by manual review in IGV, indicating that complex repair associated events were rarely recovered in standard somatic variant output from this dataset^8^. Consistent with this observation, the subsequent complex indel study noted that almost no complex indel events were evident among the common small deletion and insertion calls from Mutect2 and Strelka2, despite the expectation that nonhomologous end joining (NHEJ) repair products would frequently contain both deletion and nucleotide addition^4^. Together, these observations suggested that many complex DSB repair events were present in the data but not being preserved in a form suitable for direct interpretation. Therefore, we were motivated to develop the present simulation-based framework to detect these events.

### FreeBayes most often represented simulated complex indels as single nearby variants

A total of 2,000 complex indels were introduced across the selected chromosome 22 interval and used as the test set for comparison of variant calling workflow output. For the initial analysis, reported variants were tabulated if they occurred within 5 bases of a simulated event to account for local differences in positional representation. Under this framework, FreeBayes produced 1,986 variant calls within 5 bases of the simulated complex indels, a value close to the total number of simulated events. Of the 2,000 simulated complex indels, 1,934 were represented by exactly one nearby FreeBayes variant, corresponding to 96.7% of simulated sites (Table 1 and Fig. 1A). DRAGEN Somatic also frequently represented simulated sites as single nearby calls, with 1,762 of 2,000 simulated sites represented by exactly one nearby call. By contrast, HaplotypeCaller, Mutect2, Strelka2, and DRAGEN Germline represented 740, 767, 830, and 739 simulated sites, respectively, as exactly one nearby variant call, corresponding to 37.0%, 38.4%, 41.5%, and 37.0% of simulated sites (Table 1 and Fig. 1A). These findings indicate that FreeBayes most often preserved one-to-one representation of simulated complex indels among the workflows examined.

**Table 1.** Variant-caller representation of 2,000 simulated complex indels within a 5 bp window.

| Variant caller | Total nearby calls | Mean nearby calls per simulated site | 0 calls | 1 call | 2 calls | 3 calls | 4 calls | 5 calls |
| --- | --- | --- | --- | --- | --- | --- | --- | --- |
| FreeBayes | 1986 | 0.99 | 40 (2.0%) | 1,934 (96.7%) | 26 (1.3%) | 0 (0.0%) | 0 (0.0%) | 0 (0.0%) |
| HaplotypeCaller | 3638 | 1.82 | 40 (2.0%) | 740 (37.0%) | 826 (41.3%) | 331 (16.6%) | 62 (3.1%) | 1 (0.1%) |
| Mutect2 | 3514 | 1.76 | 42 (2.1%) | 767 (38.4%) | 856 (42.8%) | 306 (15.3%) | 28 (1.4%) | 1 (0.1%) |
| Strelka2 | 3476 | 1.74 | 66 (3.3%) | 830 (41.5%) | 695 (34.8%) | 380 (19.0%) | 29 (1.5%) | 0 (0.0%) |
| DRAGEN-germline | 3637 | 1.82 | 41 (2.1%) | 739 (37.0%) | 826 (41.3%) | 331 (16.6%) | 62 (3.1%) | 1 (0.1%) |
| DRAGEN-somatic | 2156 | 1.08 | 41 (2.1%) | 1,762 (88.1%) | 197 (9.8%) | 0 (0.0%) | 0 (0.0%) | 0 (0.0%) |

**Table note:** Percentages are calculated from the 2,000 simulated complex indel sites. “Total nearby calls” represents the total number of variant records reported within 5 bp of the simulated sites. “Mean nearby calls per simulated site” is calculated as total nearby calls divided by 2,000.

**Figure 1.**
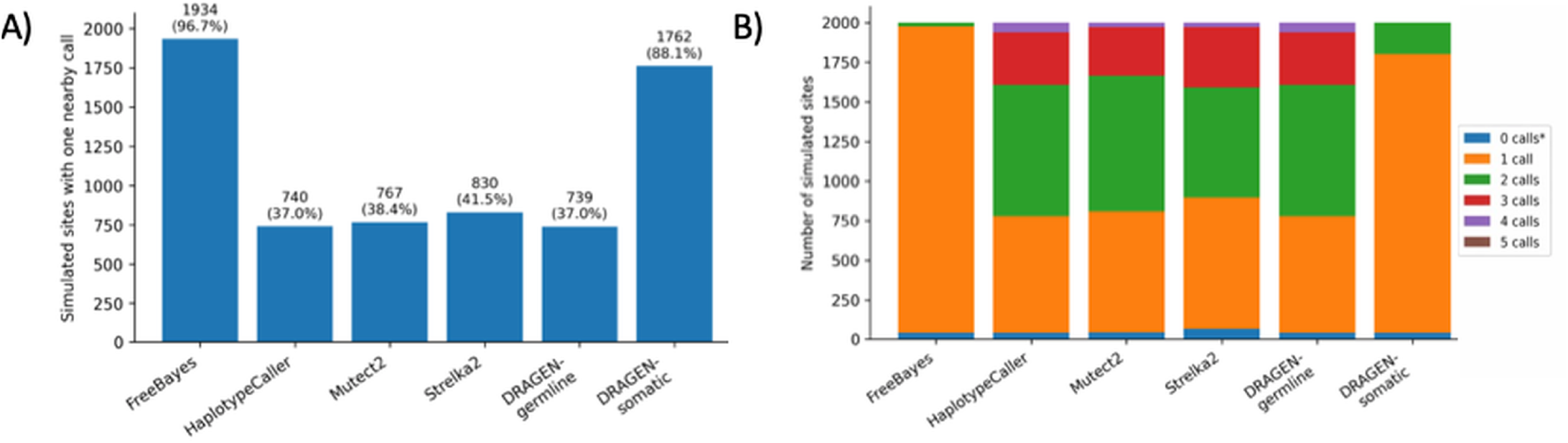
Variant-caller representation of simulated complex indels. A total of 2,000 simulated complex indels were introduced into a defined region of chromosome 22 and compared with nearby variant calls reported by FreeBaves, HaplotypeCaller, Mutect2, Strelka2, DRAGEN-germline, and DRAGEN-somatic. **A)** Number and percentage of simulated sites represented by exactly one nearby variant call. FreeBayes represented 1,934 of 2,000 simulated sites as one nearby call, compared with 740 for HaplotypeCaller, 767 for Mutect2, 830 for Strelka2, 739 for DRAGEN-germline, and 1,762 for DRAGEN-somatic. **B)** Distribution of simulated sites according to the number of nearby calls reported within the 5 bp window. FreeBayes most often preserved one-to-one representation, whereas HaplotypeCaller, Mutect2, Strelka2, and DRAGEN-germline more frequently represented a single simulated complex indel as multiple nearby calls.

### Several variant callers fragmented one complex event into multiple nearby calls

The excess number of nearby calls from several callers reflected a tendency to decompose a single simulated complex indel into multiple simpler events. HaplotypeCaller produced 3,638 nearby calls, Mutect2 produced 3,514, Strelka2 produced 3,476, and DRAGEN Germline produced 3,637 (Table 1). For HaplotypeCaller, 826 simulated sites were represented by two nearby calls, 331 by three calls, 62 by four calls, and 1 by five calls. Mutect2 showed a similar pattern, with 856 simulated sites represented by two nearby calls, 306 by three calls, 28 by four calls, and 1 by five calls. Strelka2 also showed substantial fragmentation, with 695 simulated sites represented by two nearby calls, 380 by three calls, and 29 by four calls. DRAGEN Germline closely resembled HaplotypeCaller, with 826 simulated sites represented by two nearby calls, 331 by three calls, 62 by four calls, and 1 by five calls. In contrast, FreeBayes represented only 26 simulated sites with two nearby calls and none with three or more nearby calls,while DRAGEN Somatic represented 197 simulated sites with two nearby calls and none with three or more nearby calls (Table 1 and Fig. 1B). This difference indicates that the main advantage of FreeBayes in this setting was not simply detection near the locus, but preservation of the compound event as a single interpretable record.

### A representative example illustrated complete recovery by FreeBayes and fragmentation by the other callers tested

Analysis of a representative simulated event at chromosome 22 position 30,005,501 illustrates the differences in variant representation described above. This event consisted of deletion of GGTTTGGA with insertion of TTCG, yielding a net loss of four nucleotides (Fig. 2A and 2B). FreeBayes reported the event as a single nearby variant call, which fully recapitulated the event after trimming of shared flanking bases (Fig. 2C). In contrast, HaplotypeCaller, Mutect2, and DRAGEN Germline pipeline represented the same event as three nearby variant calls rather than as one compound event (Fig. 2D). Strelka2 also represented the event as three nearby calls (Fig. 2E). DRAGEN Somatic pipeline similarly represented the event as two nearby calls rather than as a single complex-indel record (Fig. 2F). This example shows that positional recovery alone is not sufficient for complex indels. Accurate biological interpretation requires preservation of the effective deleted and inserted sequence within one coherent allele representation.

**Figure 2.**
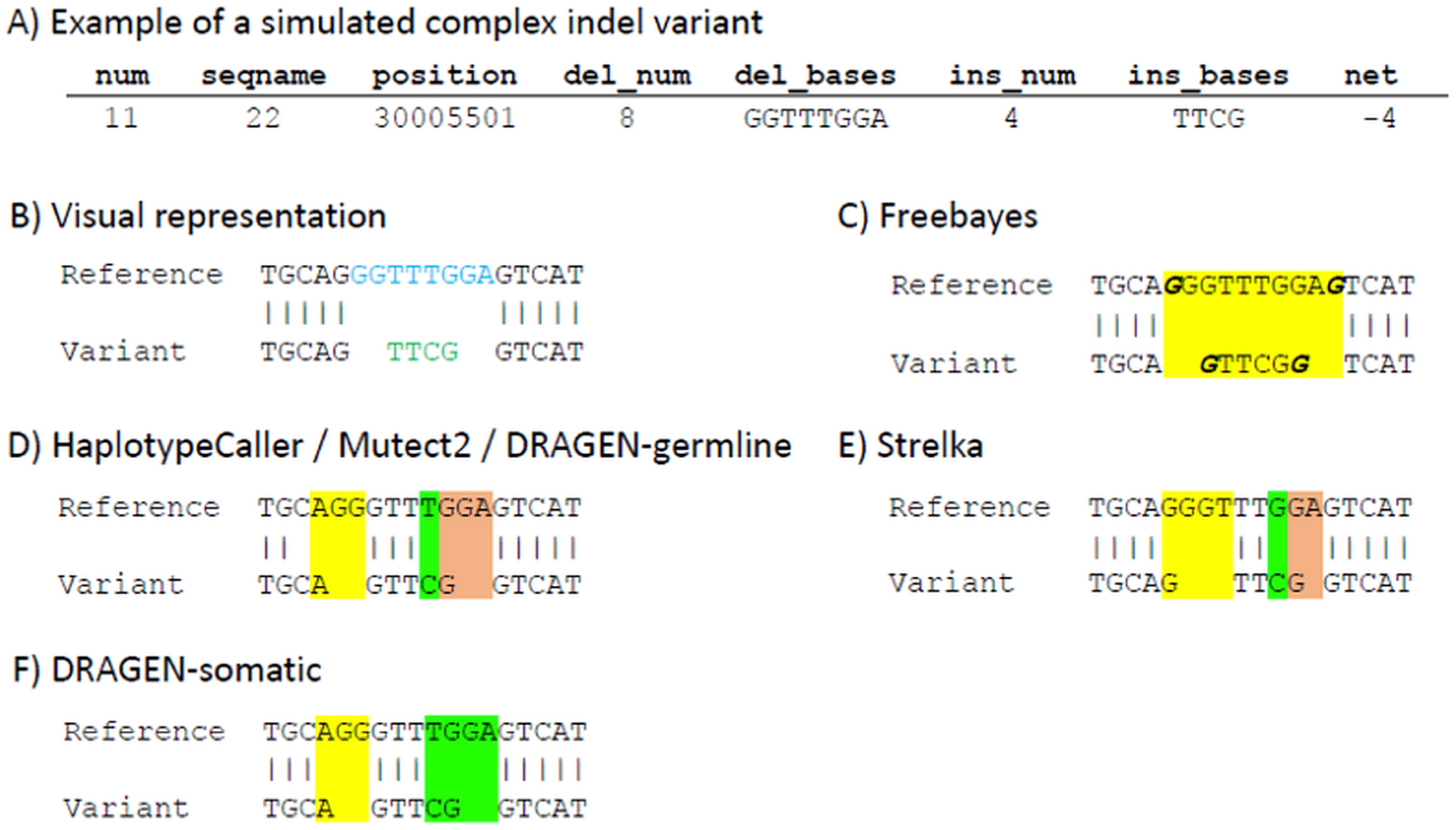
Representative complex indel call across variant callers. A simulated complex indel at chromosome 22 position 30,005,501 consisted of the deletion of GGTTTGGA and the insertion of TTCG, resulting in a net loss of four nucleotides. **A)** Simulated event record. **B)** Alignment-based representation of the simulated sequence change. **C)** FreeBayes reported the event as a single nearby variant call that retained the effective deleted and inserted sequence after trimming shared flanking bases. **D)** HaplotypeCaller, Mutect2, and DRAGEN-germline represented the same event as three nearby variant calls. **E)** Strelka2 also represented the event as three nearby calls rather than one unified complex-indel record. **F)** DRAGEN-somatic represented the event as two nearby calls rather than as a single complex-indel call. This example illustrates that positional recovery alone is insufficient for interpreting complex indela when the deleted and inserted sequences are not retained within a single variant record. Blue and green text show the simulated deletion and insertion bases respectively. Bold italicized text show shared flanking bases. Each variant record highlighted with different colors.

### Most single FreeBayes calls matched the simulated event after accounting for effective allele representation

To examine how closely single FreeBayes calls corresponded to the simulated alleles, a merged comparison table was generated for the 1,934 simulated complex indels that had exactly one nearby FreeBayes variant (Supplementary Table S1). Among these events, 202 FreeBayes calls matched the simulated deletion and insertion without additional processing. Another 1,061 calls matched the simulated event after trimming shared prefix or suffix bases from the FreeBayes REF and ALT alleles. For the remaining events, we also considered whether the simulated deletion and insertion contained shared terminal bases, because such cases can produce equivalent effective representations of the same sequence change. This comparison identified 79 calls in which the untrimmed FreeBayes alleles matched the trimmed simulated event and 485 calls in which both the FreeBayes alleles and simulated event matched after trimming. Overall, 1,827 of 1,934 one-to-one FreeBayes calls were concordant with the simulated complex indel after accounting for effective allele representation, while 107 remained unmatched or unresolved by this rule-based comparison. These findings indicate that most single FreeBayes calls preserved the essential event structure, even when the literal allele strings differed because of shared flanking sequence or local representation ambiguity.

## DISCUSSION

Complex indels, typically at DSB sites and most often repaired by NHEJ, present a difficult analytical problem. This is because the observed VCF representation does not always correspond directly to the underlying biological event. The same repair product may be shifted locally, extended with matching flanking bases, or decomposed into several neighboring calls depending on local sequence context, alignment behavior, and caller-specific haplotype representation. Therefore, interpretation of complex indels cannot rely only on the reported POS, REF, and ALT fields as literal descriptors of the event. In many instances, the more informative unit is the effective allele, by which we mean the nonshared deleted and inserted sequences that remain after identical prefix or suffix bases are trimmed from the reported REF and ALT alleles. This issue is especially important for DSB repair events, where local ambiguity can obscure the full structure of the junction and where a single biological event may otherwise appear as several simplified changes.

In the simulation framework used here, FreeBayes displayed a practical strength for this variant class. FreeBayes preserved complex indels as single nearby variant records more often than the other variant callers and pipelines evaluated. DRAGEN Somatic pipeline also performed well and showed strong one-to-one representation of simulated complex indels. However, FreeBayes produced the highest one-to-one representation in this analysis. Sequence-level comparison further revealed that most one-to-one FreeBayes calls were concordant with the simulated event after accounting for effective allele representation, supporting its use as the input for downstream parsing. FreeBayes also remained practical for downstream method development because it is open source and readily implementable. By contrast, HaplotypeCaller, Mutect2, Strelka2, and DRAGEN Germline more frequently fragmented complex indels into multiple simpler nearby calls. These findings indicate that the principal value of FreeBayes in this context is not merely sensitivity near the locus, but retention of the compound event in a form that remains biologically interpretable.

However, FreeBayes output still requires careful interpretation. Even when one nearby call corresponds to one simulated event, the reported allele often includes additional matching bases on one or both sides of the altered sequence. In other cases, coincidental sequence identity between deleted and inserted bases creates intrinsic ambiguity such that more than one formal representation is possible. Accordingly, the raw FreeBayes call should not be treated as a final biological designation without additional processing. Side-by-side comparison of simulated and FreeBayes alleles showed that literal VCF representation alone was not sufficient for standardized interpretation of complex indels. The same biological event could be reported with a slight positional offset, with shared flanking sequence appended to both alleles, or in a form influenced by local sequence ambiguity. Therefore, the effective allele emerged as the most informative interpretive unit. This strategy distinguished true complex indels from effective single-nucleotide variants, simple insertions, and simple deletions, and provided a consistent basis for downstream filtering, comparison, and mechanistic inference.

More broadly, these observations carry an important caution for the interpretation of complex indels from sequencing data in general. A variant caller that performs well for single-nucleotide variants or simple indels should not automatically be assumed to provide equally reliable representation of compound DSB repair junctions. For this event class, fragmentation into multiple nearby calls can make a true complex indel appear absent, inflate the apparent number of local events, or distort the inferred repair structure. This point is particularly relevant for studies of nonhomologous end joining, where small deletions, nucleotide addition, and nearby base changes may arise as part of one repair outcome. In such settings, failure to preserve the event as a single interpretable allele can directly limit biological inference. For complex indels, failure of biologically faithful representation is a central problem, even when the correct locus is detected.

This problem first became clear during prior analysis of the colon crypt dataset. Complex DSB repair events were biologically expected in this setting, yet they were not readily visible in standard somatic variant output from Mutect2 and Strelka2. This raised the possibility that many such events were present in the data but were not being preserved in a form suitable for direct interpretation. The simulation framework presented here was developed to examine that specific representational problem.

These observations also shaped the downstream parsing workflow. The simulation results demonstrated that FreeBayes often retained complex indels as single variant records. These results also showed that interpretation depended on the effective sequence change rather than on the unprocessed POS, REF, and ALT fields alone. Building on these observations, a rule-based workflow was developed to process FreeBayes output before manual review in IGV. This workflow excludes common and low-complexity events, derives effective deleted and inserted sequences through trimming of shared allele sequence, and applies depth, allele frequency, repeat overlap, and sequence-context filters. The simulation framework did not merely compare caller behavior. It established the interpretive logic that made a FreeBayes-centered strategy practical for analysis of complex variants in the whole-genome colon crypt dataset, in which 385 NHEJ events were detected after additional manual curation. It is also important to note that this strategy was designed to emphasize ruling out false positives; therefore, some true complex variants may be missed.

This work was not intended as a universal benchmark of all available somatic and germline variant callers. Benchmarking of somatic point mutation and indel callers has been extensively addressed in prior comparison studies and reviews^1,22–27^. The performance of variant callers varies with sequencing depth, allele frequency, dataset design, and variant class. Rather, the workflows examined here were selected because they were available within the analysis environment, commonly used in practice, and directly relevant to the representation problem observed in the colon crypt dataset. Because the simulation included both an altered test sample and an unchanged matched control sample, Mutect2, Strelka2, and DRAGEN Somatic pipeline could be run in paired tumor-normal mode, with the altered test sample treated as the tumor-equivalent sample and the unchanged sample treated as the normal-equivalent control. FreeBayes, HaplotypeCaller, and DRAGEN Germline pipeline were included because they provide useful comparisons for how germline workflows represent the same engineered complex indel events. The purpose of including these workflows was not to assign germline status, but to compare allele representation across commonly used variant calling frameworks. Simulated reads were generated with an available ART Illumina error model rather than a NovaSeq X sequencing profile, and the simulation was therefore intended to compare relative caller behavior under controlled sequencing conditions rather than to reproduce exact platform-specific error characteristics.

In conclusion, complex indel analysis depends not only on detecting a variant near the expected locus, but also on preserving the full deletion-insertion structure in a biologically interpretable form. In this simulation framework, FreeBayes most consistently retained complex indels as single compound records, while DRAGEN Somatic also showed strong one-to-one representation. HaplotypeCaller, Mutect2, Strelka2, and DRAGEN Germline more often fragmented the same underlying event into simpler neighboring calls. Because FreeBayes combined the strongest one-to-one performance with open-source availability, it was selected for downstream parsing. This two-stage strategy uses FreeBayes variant calling followed by parsing of the effective deleted and inserted sequences, enabling more consistent detection and interpretation of complex DSB repair variants as single compound events.

## ACKNOWLEDGEMENTS

This work is supported by funds from NIA R01 AG 067615 and the Catherine and Joseph Aresty Endowment to CLH and ZM. ML was supported by NIH NIGMS R35 118009.

## SUPPLEMENTAL TABLE

**Supplemental Table 1. Sequence-level comparison of simulated complex indels and corresponding FreeBayes variant calls**. The table provides the complete sequence-level comparison for the 1,934 simulated complex indels that were represented by exactly one nearby FreeBayes variant call within 5 bp of the simulated position. For each simulated event, the table lists the simulated deleted and inserted sequences, the FreeBayes-reported POS, REF, and ALT alleles, and a category assignment used for sequence-level concordance analysis. Category 1 indicates direct concordance between the FreeBayes alleles and the simulated deletion and insertion. Category 2 indicates concordance after trimming shared prefix or suffix bases from the FreeBayes REF and ALT alleles. Category 3A indicates concordance between the untrimmed FreeBayes alleles and the effective simulated event after trimming shared terminal sequence from the simulated deletion and insertion. Category 3B indicates concordance after both the FreeBayes alleles and the simulated event are reduced to their effective representations. Category 3 indicates calls that remained unmatched or unresolved by this rule-based comparison.

